# Time-course transcriptome landscape of achene development in lettuce

**DOI:** 10.1101/2020.08.03.233387

**Authors:** Chen Luo, Shenglin Wang, Kang Ning, Zijing Chen, Jingjing Yang, Yixin Wang, Meixia Qi, Qian Wang

## Abstract

Lettuce (*Lactuca sativa* L.), which belongs to the large Asteraceae (Compositae) family, breeds by sexual reproduction and produce seeds. Actually, lettuce seeds are achenes, which are defined as fruits. However, few studies have described the morphological characteristics of the lettuce achenes, and genes essential for achene development are largely unknown in lettuce. To investigate the gene activity during achene development and determine the possible mechanisms that influence achene development in lettuce, we performed a time-course transcriptome analysis of lettuce achenes. A total of 23,790 expressed genes were detected at the five achene development stages. We investigated the gene expression patterns during achene development and identified the enriched biological processes at the corresponding stages. Kyoto Encyclopedia of Genes and Genomes and Gene Ontology analyses revealed a variety of transcriptomic similarities and differentiation at different achene development stages. Further, transcription factors and phytohormones were found to play important roles during achene development. Finally, we proposed a working model to illustrate the gene expression modules and possible molecular mechanism underlying achene development. Our time-course transcriptome data also provides a foundation for future functional studies to reveal the genetic control of achene development in lettuce.

## 1. Introduction

Lettuce (*Lactuca sativa* L.) supplies dietary fiber, vitamins, and minerals, and is one of the most important leafy vegetables consumed worldwide (Zhang *et al*., 2017). Lettuce belongs to the large Asteraceae (Compositae) family (Reyes-Chin-Wo *et al*., 2017). It is a heat-promoted bolting vegetable, so the transition from vegetative growth to reproductive growth is accelerated under high temperatures (Han *et al*., 2016). Finally, the lettuce plants begin to bloom and bear fruit. Previously, we characterized the inflorescence development of lettuce (Chen *et al*., 2017), and proposed a model for lettuce floral organ specification (Ning *et al*., 2019). Lettuce seeds are achene fruits that are developed from the ovaries. However, gene activity during lettuce achene development and the general mechanisms influencing achene development are still largely unknown.

In angiosperms, the ovule develops into a seed upon fertilization, whereas the ovary differentiates into a fruit (Ruan *et al*., 2012). Many fruit types have evolved as flowering plants have adapted to different conditions (Seymour *et al*., 2013). Fruit types can be classified according to the following characteristics: carpel number, free or fused carpels, texture, dehiscence or indehiscence. Fruits derived from a mature ovary and containing seeds are defined as true fruits. However, some fruits are composed of a variety of tissues other than the ovary, such as bracts, sepals, petals, and receptacle, and these fruits are defined as false fruits. In general, fruits can be divided into two main categories: fleshy fruits and dry fruits (Dardick and Callahan, 2014). Fleshy fruits include berries, pomes, drupes, pepos, and hesperidia. Dry fruits can be either dehiscent or indehiscent depending on whether or not the pericarp splits open at maturity. In dehiscent fruits, including siliques, capsules, and legumes, the fruits open to disperse the seeds during maturation. In indehiscent fruits, including achenes, caryopses, and nuts, the fruits do not open during maturation.

The fruit of *Arabidopsis* has been considered as a model to study fruit development. The fruit is mostly made up of an ovary with three distinct tissues, the valve, the replum, and the valve margin (Roeder and Yanofsky, 2006). Until now, crucial genetic networks that regulate fruit development have been uncovered mainly in *Arabidopsis* (Pabon-Mora *et al*., 2014). For example, *FRUITFULL* (*FUL*) plays a direct role in promoting valve development (Gu *et al*., 1998); *REPLUMLESS* (*RPL*) and *BREVIPEDICELLUS* (*BP*) are necessary for replum identity (Venglat *et al*., 2002; Roeder *et al*., 2003), and *ASYMMETRIC LEAVES1* (*AS1*) and *ASYMMETRIC LEAVES2* (*AS2*) regulate replum development by repressing *BP* and *RPL* (Alonso-Cantabrana *et al*., 2007). Furthermore, *INDEHISCENT* (*IND*) and *ALCATRAZ* (*ALC*) are necessary for correct valve margin formation in fruit development (Rajani and Sundaresan, 2001; Liljegren *et al*., 2004), and *SHATTERPROOF1* (*SHP1*) and *SHATTERPROOF2* (*SHP2*) regulate valve margin formation by positively regulating *IND* and *ALC* (Liljegren *et al*., 2000). Fruit development and seed development are highly synchronized (Robert, 2019). Fruits protect the developing seeds and contribute to seed dispersal. The seed is composed of seed coat, embryo, and endosperm (Sreenivasulu and Wobus, 2013). Seed development is a complex process that is controlled by multiple biological processes and signaling pathways, such as the ubiquitin-proteasome pathway, phytohormone biosynthesis and signal transduction, and transcriptional regulators (Li and Li, 2016). Many important regulators associated with seed development and seed size determination have been identified (Li *et al*., 2019), including *DA1, DA1-related1* (*DAR1*), *AUXIN RESPONSE FACTOR2* (*ARF2*), *BRASSINOSTEROID INSENSITIVE1* (*BRI1*), *BRASSINAZOLE-RESISTANT1* (*BZR1*), *APETALA2* (*AP2*), *AINTEGUMENTA* (*ANT*), and *TRANSPARENT TESTA GLABRA2* (*TTG2*).

Many transcriptome profiling studies have been performed to elucidate the gene dynamic activity and biological processes during fruit or seed development in different plant species. In *Arabidopsis*, a time-course transcriptome analysis of the siliques identified genes that control fruit formation and maturation, and revealed the molecular mechanisms affecting silique development and maturation (Mizzotti *et al*., 2018). In kiwifruit, an integrated analysis of the fruit metabolome and transcriptome provided insights into the regulatory network of flavonoid biosynthesis during fruit development (Li *et al*., 2018). In chickpea, a global transcriptome and co-expression network analysis revealed the molecular mechanism underlying seed development, and identified candidate genes associated with seed development and seed size/weight determination (Garg *et al*., 2017). In maize, a high temporal-resolution transcriptome landscape of early maize seed development was reported. The high-density, time-course transcriptome data provided a high-resolution gene expression profile and demonstrated four key stages during early seed development in maize (Yi *et al*., 2019).

Achenes are indehiscent dry fruits of, for example, strawberry, buckwheat, and Asteraceae plants. Achenes are single-seeded fruits, and a mature achene has four major components: a hard and relatively thick pericarp, a thin testa (integument), an embryo, and an endosperm. However, few studies have analyzed the transcriptomes of achenes to explore the gene activity and molecular mechanisms involved in achene development, especially in Asteraceae plants. In this study, we identified representative developmental stages of lettuce achenes and performed time-course transcriptome analysis of the achenes at five developmental stages. We analyzed gene activity during achene development and proposed a possible mechanism that may influence achene development. Our time-course transcriptome data provide an important resource for future functional studies and shed light on the genetic control of achene development in lettuce.

## 2. Materials and Methods

### 2.1 Plant materials and growth conditions

The lettuce cultivar S39 was used in this study. In the vegetative growth stage, the lettuce plants were grown in growth chambers under a cycle of 16-h light at 25°C and 8-h dark at 18°C. When the lettuce plants transitioned to reproductive growth, they were transplanted to a greenhouse under standard conditions.

### 2.2 Sample collection and RNA extraction

Achene samples were collected at the S0, S1, S3, S6, and S9 stages for RNA-sequencing (RNA-seq). The achenes were harvested from at least five capitulum for each pooled sample. A total of 15 samples, from five developmental stages with three biological replicates, were collected and stored in an ultra-low temperature freeze at –80°C. Total RNA was extracted from each sample using a Quick RNA isolation kit (Huayueyang, Beijing, China) according to the manufacturer’s protocol.

### 2.3 RNA-seq and bioinformatic analysis

Paired-end libraries were constructed and sequenced using an Illumina Nova-PE150 platform at Novogene (Beijing, China). Bioinformatic analysis of the RNA-seq data was performed on the BMKCloud platform (www.biocloud.net). Raw reads were filtered to generate the clean reads, and the clean reads were mapped to the lettuce reference genome using the HISAT2 software (Kim *et al*., 2015). Transcript abundances were determined by the fragments per kilobase of transcript per million mapped reads (FPKM) values, and genes with FPKM values ≥0.1 were retained for further analysis. Principle component analysis (PCA) was performed using the prcomp function in R software (R Core Team, 2014).

### 2.4 Gene expression pattern analysis

To further explore the gene expression patterns during achene development, clustering analysis was performed using the K-means method with the Pearson correlation distance in the MeV software (https://sourceforge.net/projects/mev-tm4/). The expression level of each gene was normalized by the average FPKM values of the different achene development stages. Then, 12 clusters were generated, and the heatmaps for each cluster were created using the normalized FPKM values of the different achene development stages.

### 2.5 Functional annotation

Genes were functionally annotated with the NCBI non-redundant protein sequences database using the BLAST software. Gene Ontology (GO) enrichment analysis was performed using the GOseq R packages (Young *et al*., 2010). GO enrichment heatmap for the genes in five of the clusters (Cluster1–Cluster5) was created using the TBtools software (Chen *et al*., 2020). Kyoto Encyclopedia of Genes and Genomes (KEGG) analysis was performed using the KOBAS software (Mao *et al*., 2005).

## 3. Results

### 3.1 Representative achene development stages of lettuce

The lettuce inflorescence is a capitulum that consists of multiple ray florets, each of which has five major components: petal, pappus, stamen, pistil, and ovary (Fig. 1A). The ovary develops into an achene, often referred to as the seed in lettuce. We defined the achene development stages according to the number of days after pollination, and the representative stages are as follows: S0 stage, the day of pollination (the processes of pollination and double fertilization take place, and the achene begins to develop); S1 stage, 1 day after pollination (the achene elongates rapidly); S3 stage, 3 days after pollination (the achene grows longer and wider); S6 stage, 6 days after pollination (the achene expands and obvious longitudinal ribs appear on the achene surface); and S9 stage, 9 days after pollination (the longitudinal ribs are more pronounced and the achene coat becomes darker, and the achene is fully developed and ready to ripen) (Fig. 1B).

**Fig. 1.**
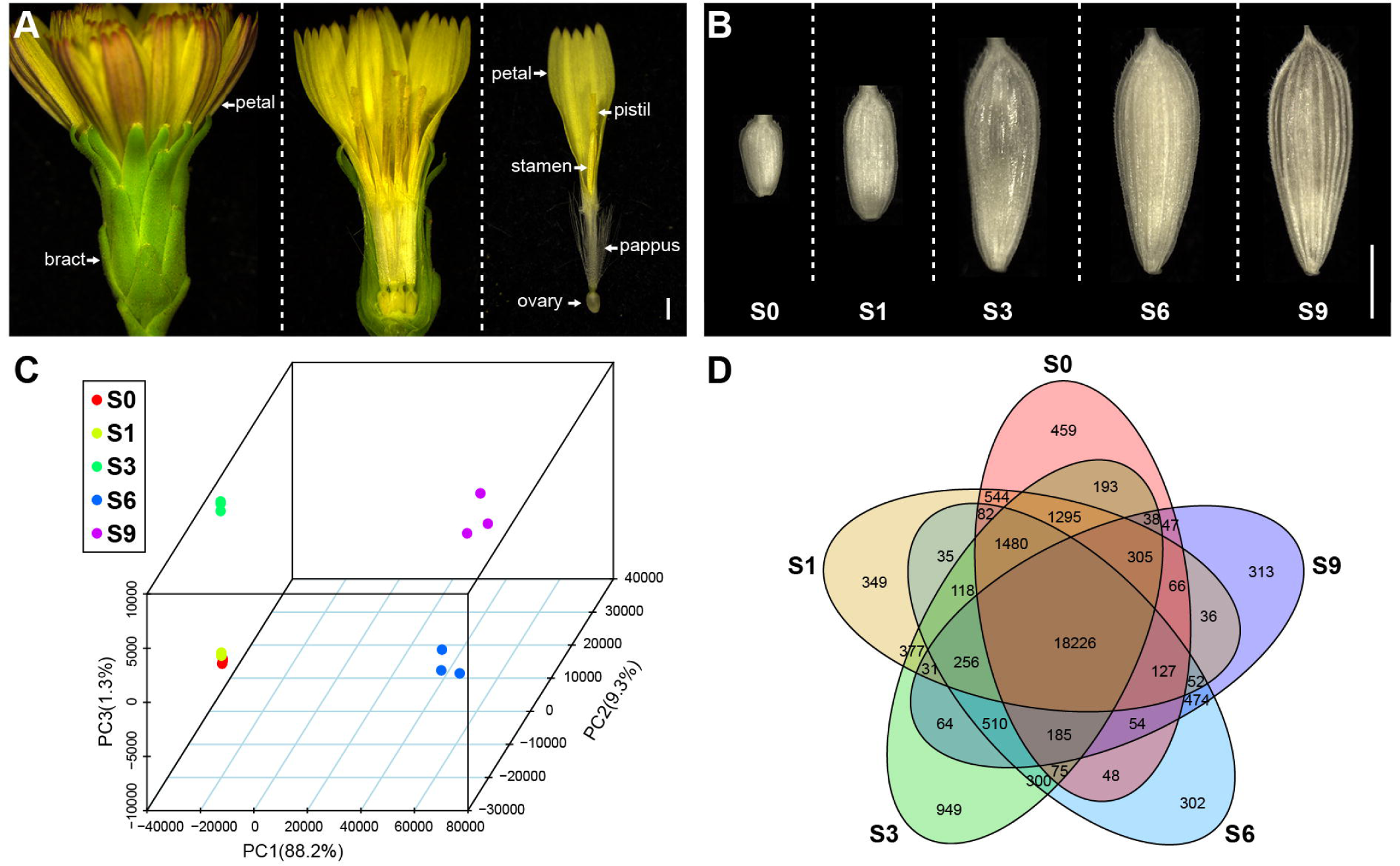
Morphological characterization of the lettuce achenes and global analysis of the transcriptome data. (A) Morphological characterization of the lettuce capitulum and floret. (B) Morphology of achenes at different developmental stages. (C) PCA of gene expression in different achene samples. (D) Venn diagram for gene expression analysis in different achene samples. Scale bars, 1 mm.

### 3.2 Global analysis of the time-course transcriptome data from different achene samples

To investigate the gene activity and analyze the molecular networks during achene development in lettuce, we performed time-course transcriptome analysis of the achenes from the five developmental stages, with three biological replicates for each time point. A total of 730.58 million clean reads (109.59 Gb clean bases) were generated from the 15 samples, with 39.31–57.45 million clean reads for each sample (Table S1). Clean reads were mapped to the lettuce genome, and the unique mapped reads ratios were 85.60%–93.08% (Table S1).

The principal component analysis (PCA) of the transcriptome datasets showed that the 15 time-series samples formed four groups: group1 (S0 and S1), group2 (S3), group3 (S6), group4 (S9) (Fig. 1C). The S0 and S1 stages were closer than the other adjacent stages, indicating that the gene expression profiles for S0 and S1 were more similar than those for the other stages. Transcript abundances were determined by FPKM values (Fig. S1), and a total of 27,390 transcripts (FPKM >0.1) were found among the five stages (Table S2). Among them, 18,226 genes were widely expressed at the five developmental stages (Fig. 1D). A further 459, 349, 949, 302, and 313 genes were specially expressed at the S0, S1, S3, S6, and S9 stages, respectively (Fig. 1D).

### 3.3 Gene expression during different achene development stages

To investigate the gene expression patterns during achene development, we performed K-means clustering analysis for the 27,390 expressed genes and generated 12 co-expression clusters (Fig. 2A). Genes grouped into the same cluster showed similar expression patterns. The genes in five of the clusters (Cluster1–Cluster5) were mainly highly expressed at only one of the five developmental stages (Fig. 2B), indicating that they might have specific functions at the corresponding stages. These genes were functionally annotated by GO enrichment analysis (Fig. 2C).

**Fig. 2.**
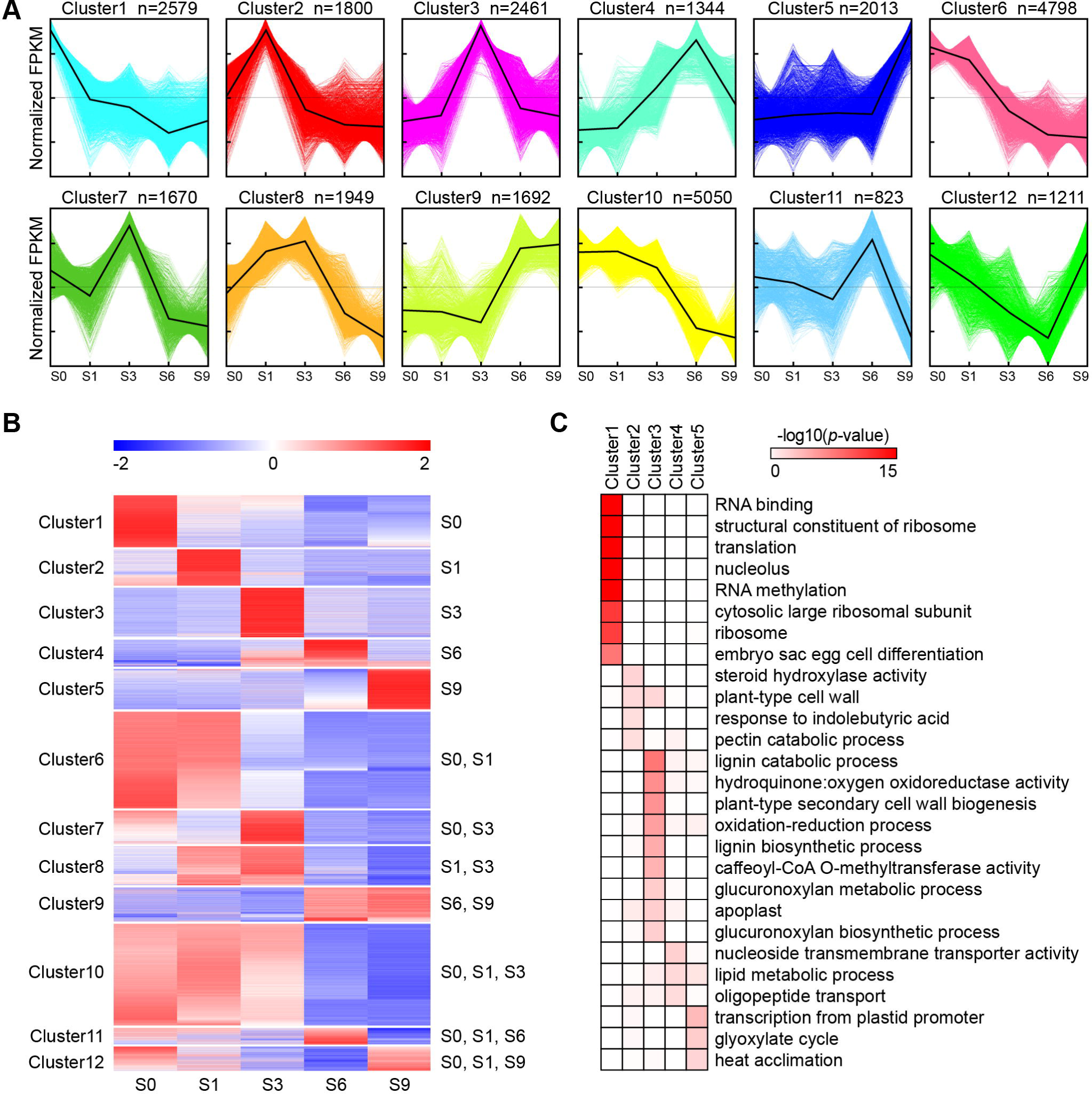
Gene expression patterns and functional transition at different achene development stages. (A) Twelve clusters for the gene expression patterns based on the normalized FPKM values. (B) Heatmaps for the gene expression patterns in different co-expression clusters. (C) GO functional enrichment analysis of the genes in different co-expression clusters.

Genes in Cluster1 were highly expressed at the S0 stage, but their expression decreased sharply at the later stages. Most of these genes were related to basic functional processes, such as RNA binding (GO:0003723), structural constituent of ribosome (GO:0003735), and translation (GO:0006412). Moreover, genes related to embryo sac egg cell differentiation (GO:0009560) also were abundant in Cluster1. S0 stage is best represented by Cluster1 because pollination and fertilization take place in this stage, which also mark the beginning of achene development. After double fertilization, cell division and cell differentiation occur more frequently and biosynthesis processes proceed rapidly. Genes in Cluster2 were more highly expressed at the S1 stage than at the other stages. Some of these genes were related to cell wall, such as plant-type cell wall (GO:0009505) and pectin catabolic process (GO:0045490), indicating that they might be associated with primary cell wall formation during achene development. Genes in Cluster3 were highly expressed only at the S3 stage. Genes related to the anabolic processes of lignin and glucuronoxylan were abundant, and genes related to plant-type secondary cell wall biogenesis (GO:0009834) were enriched in Cluster3. Glucuronoxylan is the main component of hemi-cellulose, and secondary cell walls are composed of cellulose, hemi-cellulose, lignin, pectin, and some proteins (Zhong *et al*., 2019). Because hardening of the pericarp occurs through secondary cell wall formation and lignification (Dardick and Callahan, 2014), reflecting that these enriched genes might be involved in pericarp hardening of lettuce achene. Genes in Cluster4 were more highly expressed at the S6 stage than at the other stages. These genes were related to nucleoside transmembrane transporter activity (GO:0005337), lipid metabolic process (GO:0006629), and oligopeptide transport (GO:0006857). Lipids include fatty acids and fats that are essential for energy storage and supply (Lim *et al*., 2017). The high abundance of genes associated with lipid metabolic process indicated that biochemical reactions and pathways involving lipids were more active at the S6 stage. Genes in Cluster5 had low expression at the early stages, but their expression increased significantly at the S9 stage. Some of these genes were related to transcription from plastid promoter (GO:0042793), glyoxylate cycle (GO:0006097), and heat acclimation (GO:0010286), indicating that they may be involved in plastid transcription, energy and substance metabolism, and response to high temperatures during achene development.

The genes in Cluster6–Cluster12 were expressed at more than one of the five developmental stages, which indicated that common functional processes took place at the different developmental stages. For example, Cluster10 is best presented by 5050 genes, which displayed high expression levels at the S0, S1, and S3 stages. These genes were enriched in many basic functional processes (Fig. S2), including proteasome core complex assembly (GO:0080129), response to misfolded protein (GO:0051788), and intracellular protein transport (GO:0006886).

### 3.4 Transcriptomic similarities at the different achene development stages

To investigate the transcriptomic similarities at the five developmental stages, we performed KEGG analysis of the 18,226 widely expressed genes (Fig. 3A). The most significantly enriched pathways were ribosome (ko03010), mRNA surveillance pathway (ko03015), and RNA transport (ko03013), which are involved in mRNA translation to a polypeptide chain, indicating that protein biosynthesis played a dominant role during achene development. Other pathways involved in the biosynthesis and metabolism of other substances also were active during achene development. These results indicate that biosynthesis and metabolism processes are highly active to ensure the normal development of achenes.

**Fig. 3.**
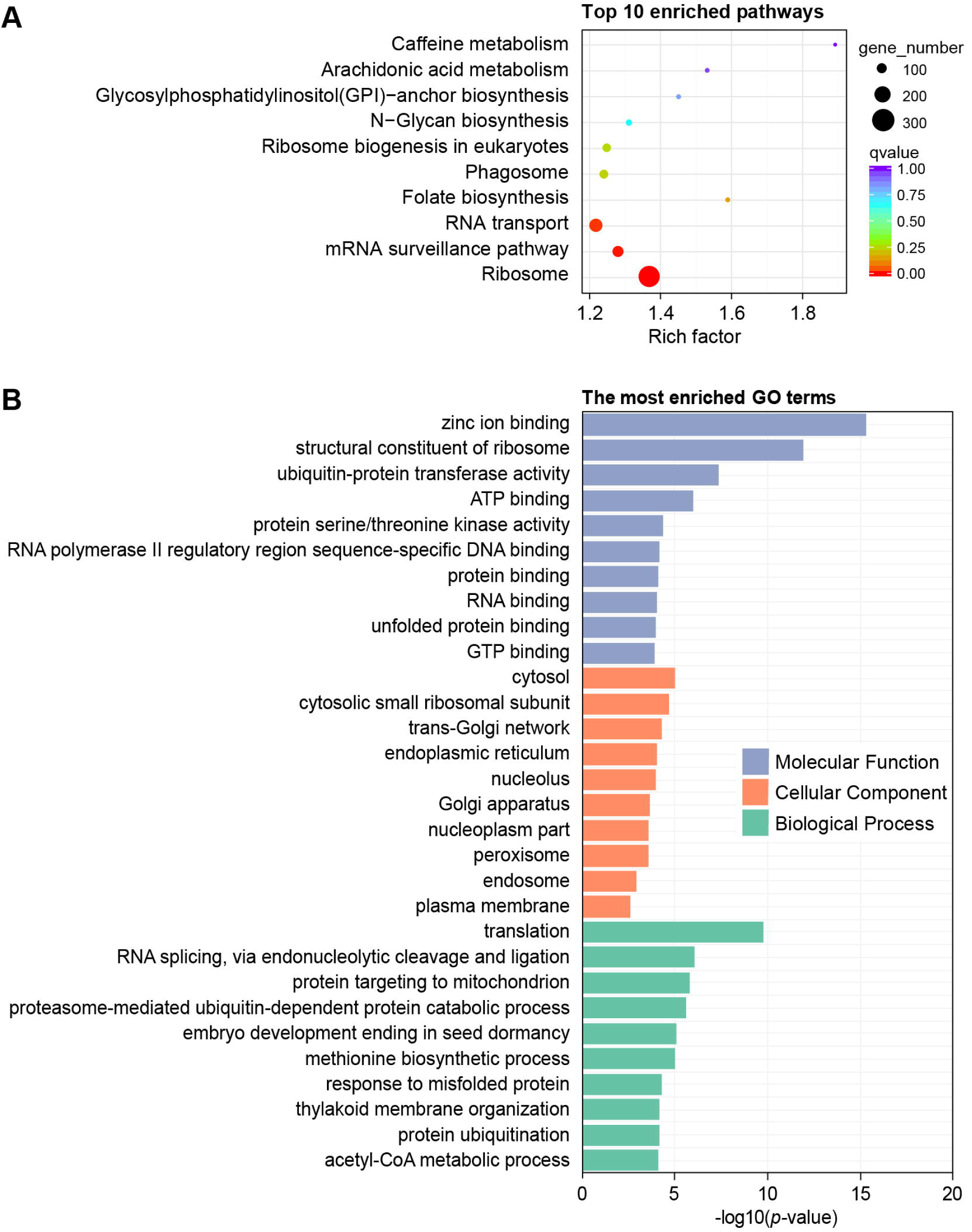
KEGG and GO analysis of the widely expressed genes at different achene development stages. (A) KEGG enrichment analysis of the widely expressed genes. (B) Significantly enriched GO terms of the widely expressed genes.

The 18,266 widely expressed genes also were assigned terms under the three main GO categories (molecular function, cellular component, and biological process) (Fig. 3B). Under molecular function, the most enriched GO terms were zinc ion binding (GO:0008270) and structural constituent of ribosome (GO:0003735). Under cellular component, the most enriched terms were cytosol (GO:0005829) and cytosolic small ribosomal subunit (GO:0022627). Under biological process, the most enriched terms were translation (GO:0006412) and RNA splicing, via endonucleolytic cleavage and ligation (GO:0000394). These genes, which were expressed at all five developmental stages, were enriched in many basic functional processes, and the results of the GO and KEGG analyses were generally consistent.

### 3.5 Transcriptomic differentiation at the different achene development stages

We analyzed the transcriptome profiling data to identify the specially expressed genes at particular stages of achene development. Then, we performed KEGG analysis of these genes in each developmental stage. At the S0 stage, the most enriched pathways were plant-pathogen interaction (ko04626) and galactose metabolism (ko00052) (Fig. 4A), which are related to environmental adaptation and carbohydrate metabolism. At the S1 stage, the most enriched pathways were phenylpropanoid biosynthesis (ko00940) and carbon fixation in photosynthetic organisms (ko00710) (Fig. 4B). Phenylpropanoids are plant secondary metabolites that function as structural and signaling molecules (Liu *et al*., 2015). Carbon fixation is essential for energy metabolism during achene development. Brassinosteroid (BR) biosynthesis (ko00905) also was active at the S1 stage (Fig. 4B), indicating that BR is an important regulator during achene development in lettuce. Genes related to flavonoid biosynthesis (ko00941) were more abundant at the S3 stage (Fig. 4C). Flavonoids are major secondary metabolites that are related to fruit flavor and color, and provide resistance against diseases and pests (Xu *et al*., 2015). At the S6 stage, biosynthesis of unsaturated fatty acids (ko01040) and fatty acid metabolism (ko01212) were the most enriched pathways (Fig. 4D). Fatty acids have multitude functions including energy storage and heat supply (Lim *et al*., 2017), which indicated that the rapid expansion of achene at this stage was accompanied with the biosynthesis and metabolism of fatty acids. Anthocyanins are common plant pigments (Jaakola, 2013), so the active anthocyanin biosynthesis (ko00942) process at the S9 stage was consistent with the color change of achene coat at this stage (Fig. 4E).

**Fig. 4.**
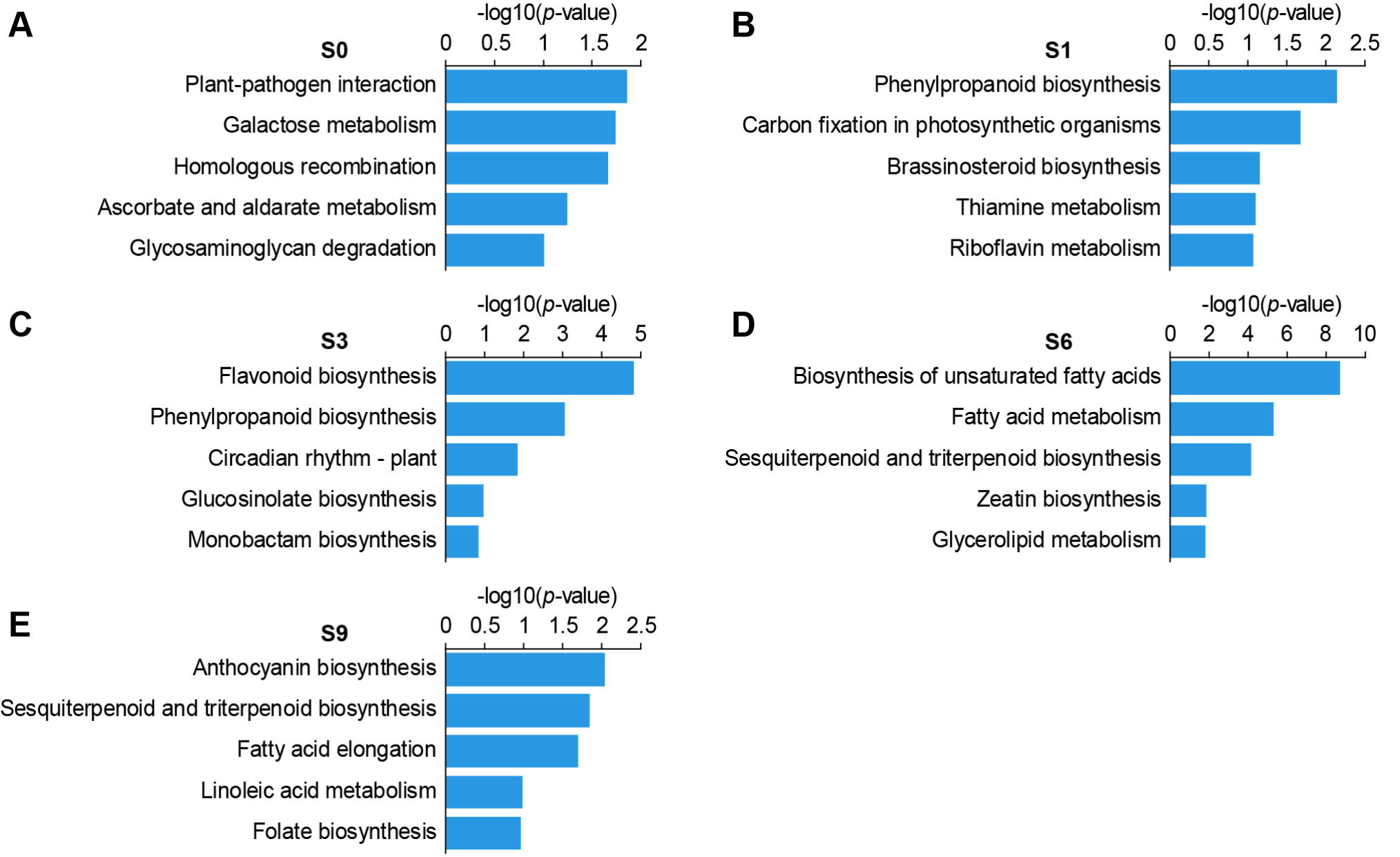
Analysis of the specially expressed genes at different achene development stages. (A–E) KEGG enrichment analysis of the stage-specific genes at the S0 (A), S1 (B), S3 (C), S6 (D), and S9 (E) stages.

### 3.6 Identification of transcription factors during different achene development stages

Transcription factors (TFs) are important regulators of transcription and gene expression, and TFs play dominant roles in regulating the growth, development, and stress tolerance of plants (Riechmann *et al*., 2000; Samad *et al*., 2017). Members of many TF families have been shown to play a wide variety of roles in controlling fruit and seed development (Sun *et al*., 2010). In this study, we identified 1579 genes that encode TFs from the transcriptome data (Table S3). Most of these TFs were expressed at all five achene development stages, but were strongly expressed at the early stages, especially at the S1 stage, although a small number of genes were highly expressed at the later stages (Fig. 5A). The top 100 highly expressed TFs at each developmental stage were classified and counted. The significantly enriched TFs mainly belonged to the MYB, APETALA2/ETHYLENE RESPONSIVE FACTOR (AP2/ERF), Homeobox (HB), MADS-box, basic leucine zipper (bZIP), and NAM/ATAF/CUC (NAC) families (Fig. 5B).

**Fig. 5.**
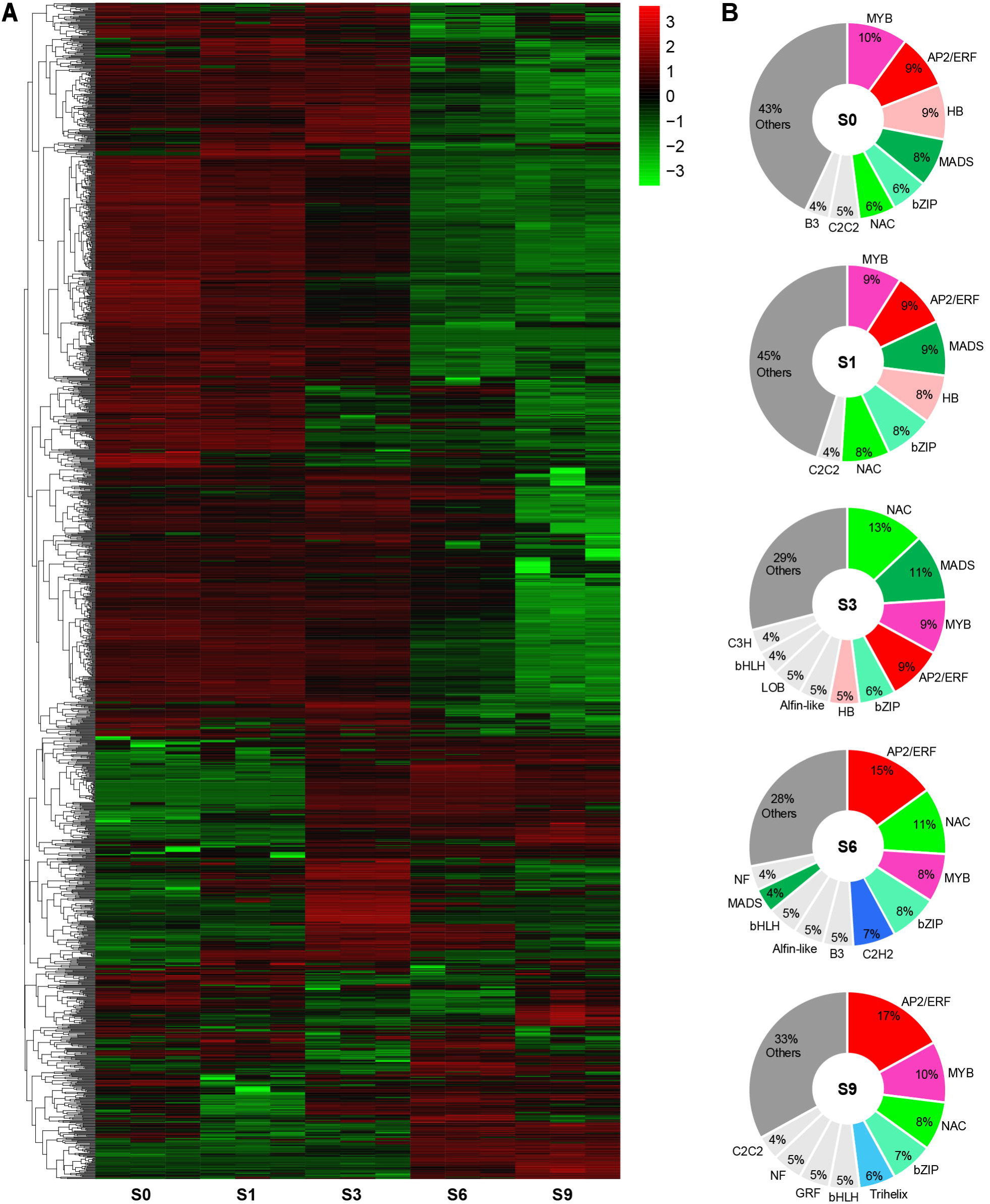
Identification of transcription factors at different achene development stages. (A) Heatmap expression profile of the identified TFs. (B) Pie charts of the top 100 highly expressed TFs at each developmental stage.

MYB TFs play prominent roles in controlling development, signal transduction pathways, the biosynthesis of secondary metabolites, and response to biotic and abiotic stresses (Dubos *et al*., 2010). We found that the MYB TFs accounted for 8%–10% of the top 100 TFs at all five developmental stages, indicating that MYB TFs functioned throughout the achene development process. The AP2/ERF TFs constitute one of the largest TF families in plants, and orchestrate the developmental processes and stress responses (Licausi *et al*., 2013; Xie *et al*., 2019). Here, we found that the AP2/ERF TFs accounted for about 9% of the top 100 TFs at the early stages, and 15% and 17% at the S6 and S9 stages respectively, reflecting the important role of AP2/ERF TFs in achene development. HB TFs, which belong to several subfamilies, regulate a large number of developmental processes in plants, including embryonic development, meristem maintenance, and organ formation (Bürglin and Affolter, 2015). Members of MADS-box TF family participate in numerous developmental process in plants, and their role in floral organ identity determination is conserved among divergent plant species. MADS-box TFs also play important roles in inflorescence architecture and fruit and seed development (Ng and Yanofsky, 2001; Smaczniak *et al*., 2012). In this study, we found that there were more HB and MADS-box TFs highly expressed at the early stages than at the later stages, suggesting that they played crucial roles at the early stages of achene development. The bZIP TFs are involved in diverse biological processes, including environmental signaling, seed maturation, flower development, and stress response (Jakoby *et al*., 2002; Droge-Laser *et al*., 2018). Here, the bZIP TFs accounted for 6%–8% of the top 100 TFs at the different developmental stages. Previous studies have revealed that NAC TFs function in developmental programs, response to environmental signals, and defense (Olsen *et al*., 2005; Puranik *et al*., 2012; Kim *et al*., 2016). Here, we found that the NAC TFs were expressed throughout the five developmental stages but had high proportions at the S3 and S6 stages, indicating that NAC TFs might have important functions at these stages. Further, the C2H2 and Trihelix TFs accounted for 7% and 6% only at the S6 and S9 stages, respectively.

### 3.7 Expression pattern of key genes associated with fruit and seed development across the achene development stages

The lettuce achene is developed from the ovary wall and ovule, thus the achene development combines both fruit development and seed development. Here, we first identified genes associated with fruit development across the development stages (Fig. 6A). Notably, the majority of these regulators are TFs. Most of the genes involved in fruit development were highly expressed at the early stages, although some genes were expressed at low levels at all five stages. We found that the lettuce homologs of *AGAMOUS* (*AG*) and *SEEDSTICK* (*STK*), which belong to the MADS-box TF family (Favaro *et al*., 2003), were up-regulated at the S0, S1 and S3 stages, indicating that they might be important for carpel development in lettuce. The *APETALA1*/*FRUITFULL* (*AP1*/*FUL*) clade of lettuce had multiple gene duplications, implying that these genes may display functional redundancy in valve development. In *Arabidopsis, SPATULA* (*SPT*) plays a minor role in promoting separation layer formation of the fruit (Heisler *et al*., 2001; Groszmann *et al*., 2011). We identified a lettuce homolog of *SPT* that showed increased expression at the S3 stage then decreased expression at the later stages. Furthermore, the expression levels of *BP* and *AS1* were higher than those of other genes associated with replum development, indicating that *BP* and *AS1* may contribute most to replum formation. Members of the KANADI (KAN) and YABBY (YAB) TF families that play central roles in lateral organ development also are crucial for ovule development in *Arabidopsis* (Kelley and Gasser, 2009). Several lettuce homologs of *KANs* and *YABs* were identified, and a *YAB1* gene (*Lsat_1_v5_gn_7_9041*) that was highly expressed across the five developmental stages might play a major role in ovule development.

**Fig. 6.**
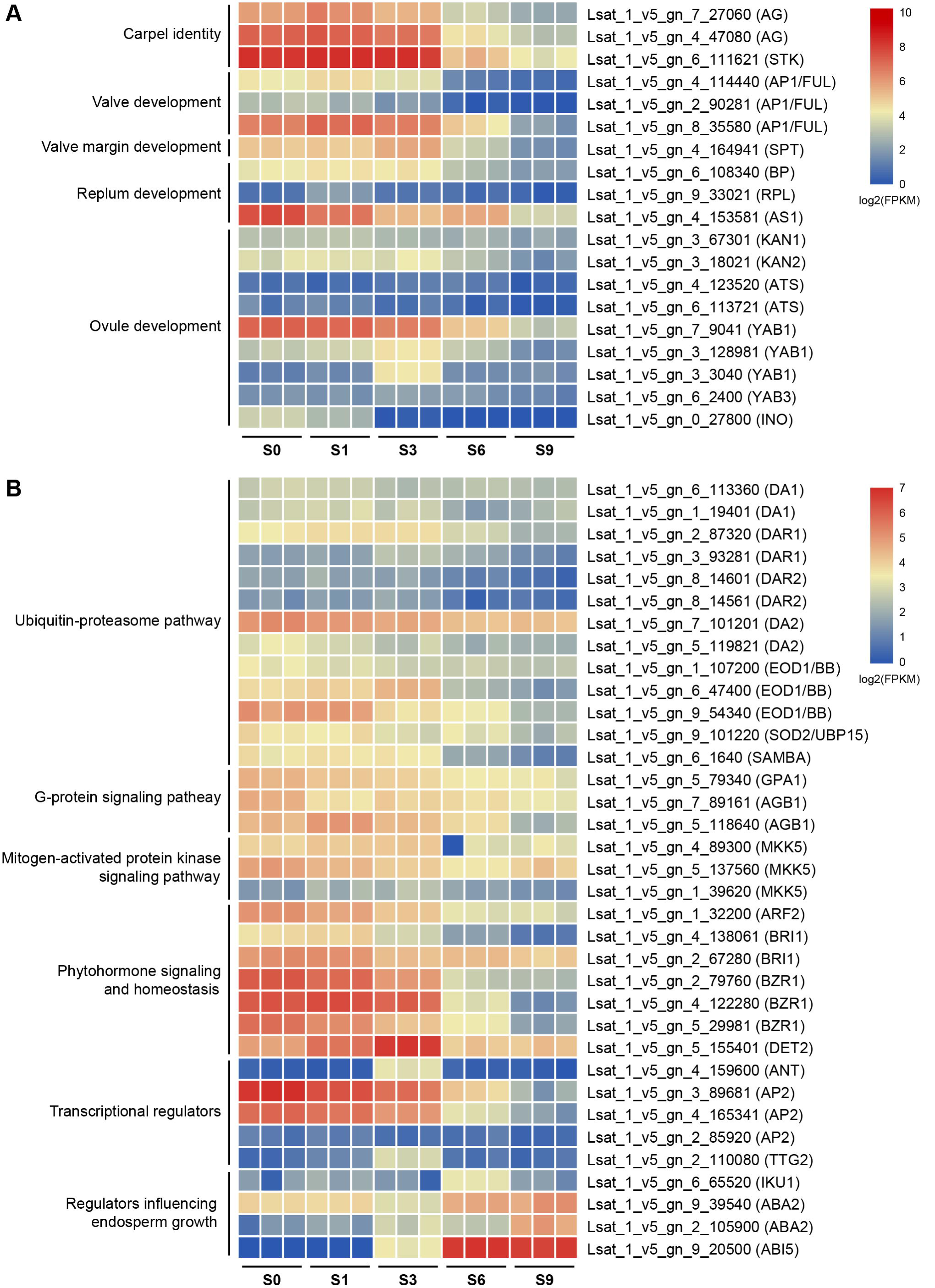
Expression patterns of key genes associated with fruit and seed development at different achene development stages. (A) Heatmap expression profile of the fruit development related genes. (B) Heatmap expression profile of the seed development related genes.

Seed development is a complex process, and many pathways involved in seed development have been proposed (Li *et al*., 2019). In this study, we identified genes associated with seed development across the five developmental stages (Fig. 6B). Genes related to the ubiquitin-proteasome pathway, G-protein signaling pathway, and the mitogen-activated protein kinase signaling pathway were stably expressed with normal expression levels at all stages. The genes related to the ubiquitin-proteasome pathway had duplicate copies, which may have resulted in functional redundancy among them. Genes related to phytohormone signaling and homeostasis and transcriptional regulators were the most highly expressed genes at the early stages, and the expression of most of them gradually decreased at the later stages. Previous studies have shown that plant hormones regulate seed growth (Sun *et al*., 2010), and the BR receptor BRI1 and the BR response TF BZR1, which are involved in BR signal transduction, were shown to control seed development (He *et al*., 2005; Kim *et al*., 2009; Huang *et al*., 2013). Here, we found that the lettuce homologs of *BRI1* and *BZR1* were highly expressed at the S0, S1, and S3 stages, and their expression decreased at the later stages. Besides, a homolog of *DE-ETIOLATED2* (*DET2*), which is involved in BR biosynthesis (Fujioka *et al*., 1997; Jiang *et al*., 2013), was more abundant at the S3 stage. These results revealed the importance of BR in regulating achene development in lettuce. The expression patterns of lettuce AP2 TFs were similar to those of the lettuce BZR1 TFs. Homologs of *ANT* and *TTG2* displayed higher expression at the S3 stage than at the other stages, indicating that *ANT* and *TTG2* may have important functions at the S3 stage. Further, genes that influence endosperm growth were highly expressed at the later stages of achene development, especially the abscisic acid (ABA) response TF ABA-INSENSITIVE5 (ABI5), which was highly expressed at the S6 and S9 stages. ABA activates seed dormancy and suppresses germination (Cutler *et al*., 2010), which indicated that the lettuce achenes were about to ripen and become dormant at the later stages.

## 4. Discussion

The lettuce seed is an achene fruit, and the development of a lettuce achene from the small ovary to its final size must involve extensive biological processes. In this study, we performed a time-course transcriptome analysis of lettuce achenes and revealed a possible mechanism that influenced achene development. To illustrate this idea, we proposed a working model of lettuce achene development (Fig. 7).

**Fig. 7.**
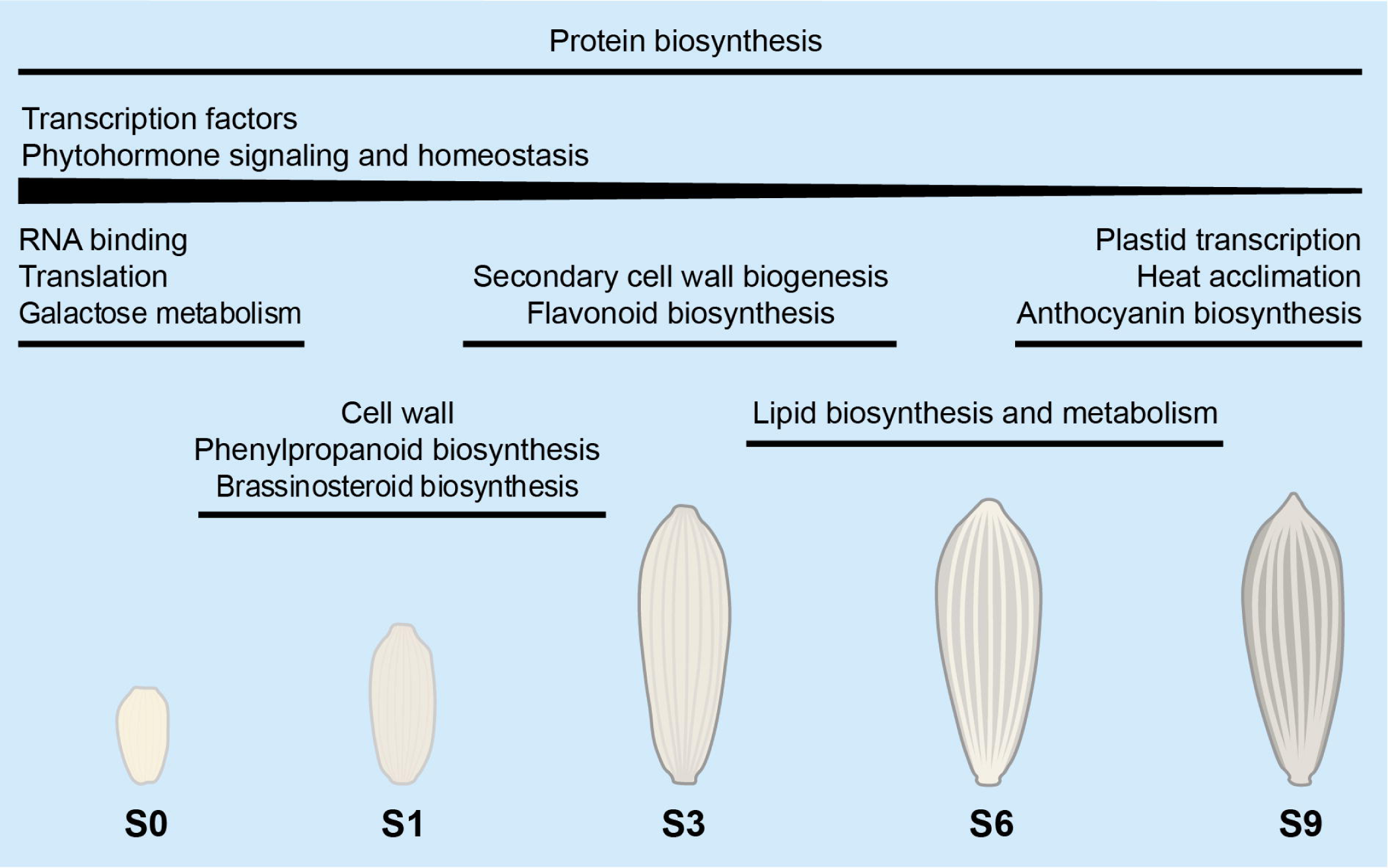
Proposed working model of achene development in lettuce. The gene expression modules and possible molecular mechanism underlying achene development in lettuce are illustrated. Biological processes and components above the lines indicate that the related genes are expressed during the corresponding achene development stages.

Overall, genes related to basic functional categories were abundant at almost every achene development stage, indicating that the basic biosynthesis and metabolism processes proceeded throughout the achene development, especially pathways associated with protein biosynthesis. These continuous and stable biological processes likely ensure the normal development of the achenes. We also found that genes that encode TFs and genes associated with phytohormone signaling and homeostasis play crucial roles during achene development. Most of these genes were highly expressed at the early stages of achene development, and their expression gradually decreased with the development of the achene.

The K-means clustering analysis identified the highly expressed genes at each developmental stage. Differential expression analysis identified genes that were particularly expressed at each stage. We also found that many biological processes, especially synthetic and metabolic activities, were significantly different at different developmental stages. At the S0 stage, the enriched pathways were RNA binding and translation, and genes related to galactose metabolism also were abundant in this stage, which reflected the rapid growth of achene after pollination and fertilization. At the S1 stage, genes related to cell wall and phenylpropanoid biosynthesis were enriched. The BR biosynthesis pathway also was active at the S1 stage, indicating the important role of BR during achene development. At the S3 stage, genes associated with the components and biogenesis process of the secondary cell wall were highly expressed, indicating that these genes may be involved in pericarp hardening of lettuce achene. The flavonoid biosynthesis pathway also was enriched at the S3 stage. Flavonoids are involved in multiple physiological functions, and many types of flavonoids in seeds are found such as flavonols and anthocyanins (Lepiniec *et al*., 2006), reflecting the important roles of flavonoids during achene development. At the S6 stage, the anabolic processes of lipids occurred more frequently, including fatty acid biosynthesis and metabolism. Lipids are essential for energy storage and supply (Lim *et al*., 2017), indicating that the rapid expansion of achene was accompanied by the accumulation of energy substances at the S6 stage. At the S9 stage, the enriched pathways included plastid transcription and heat acclimation, and genes related to anthocyanin biosynthesis also were abundant. Anthocyanins are common plant pigments and temperatures affect the stability of anthocyanins (Marszałek *et al*., 2017), so the high expression of these genes is consistent with the dark achene coat that develops at this stage.

Direct orthologs of *IND, ALC, SHP1*, and *SHP2* have not been detected in lettuce, which coincide with the report that these genes may be found only in Brassicaceae (Pabon-Mora *et al*., 2014). The fruit structure of lettuce is different from that of *Arabidopsis*; the achenes of lettuce are indehiscent fruits, whereas the siliques of *Arabidopsis* are dehiscent fruits. The four genes, *IND, ALC, SHP1*, and *SHP2*, are required for the valve margin formation and dehiscence, but the achenes do not undergo dehiscence when the fruits are fully mature. The structural differences between the fruits of lettuce and *Arabidopsis* might explain the absence of genes associated with fruit valve margin formation in lettuce.

Taken together, our data provide a foundation for future functional studies. Many genes that were highly expressed in the achenes, such as *AG, STK, BZR1, AP2*, and *ABI5*, may be responsible for important traits in lettuce, such as achene size, achene shape, grain weight, seed shattering, seed dispersal, and oil concentration. The biological functions of these genes need to be further confirmed in lettuce. The lettuce genome has been sequenced and CRISPR/Cas9-mediated genome editing has been applied to lettuce (Woo *et al*., 2015; Reyes-Chin-Wo *et al*., 2017; Zhang *et al*., 2018), so making good use of the genomics and biotechnologies will be helpful to reveal novel genetic and epigenetic variations in lettuce that can be applied to breed improved lettuce varieties in the future.

## Acknowledgments

This work was supported by Beijing Leafy Vegetables Innovation Team of Modern Agro-industry Technology Research System (BAIC07-2020) and The Construction of Beijing Science and Technology Innovation and Service Capacity in Top Subjects (CEFF-PXM2019_014207_000032).

## Supplemental Materials

Fig. S1. Boxplot for the gene expression level distribution.

Fig. S2. GO enrichment analysis of the genes in Cluster10.

Table S1. Summary of the sequencing data.

Table S2. List of genes analyzed in this study.

Table S3. Expression patterns of identified transcription factors.

