## Supplementary figures and images for "Time-course transcriptome landscape of achene development in lettuce"

### Fig. S1.tif

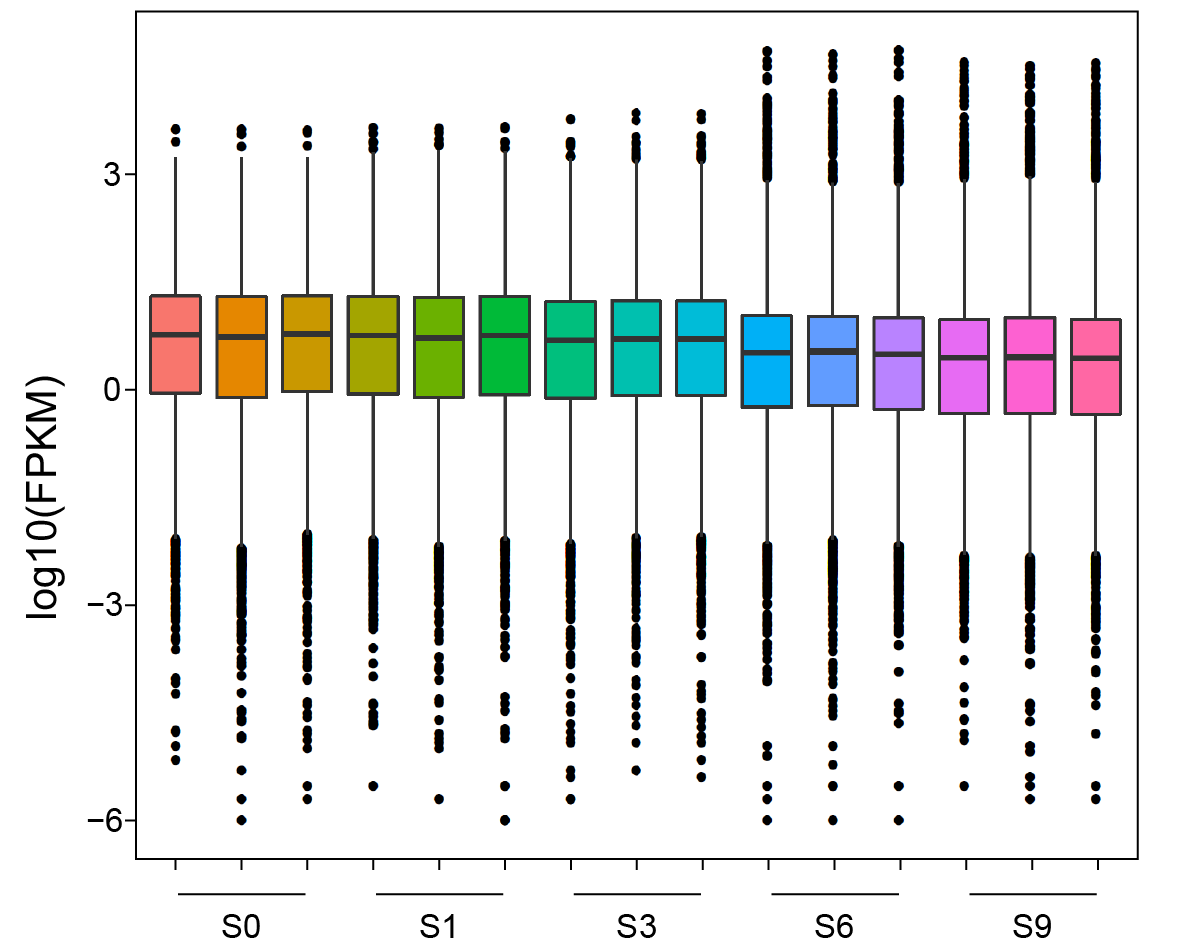

### Fig. S2.tif

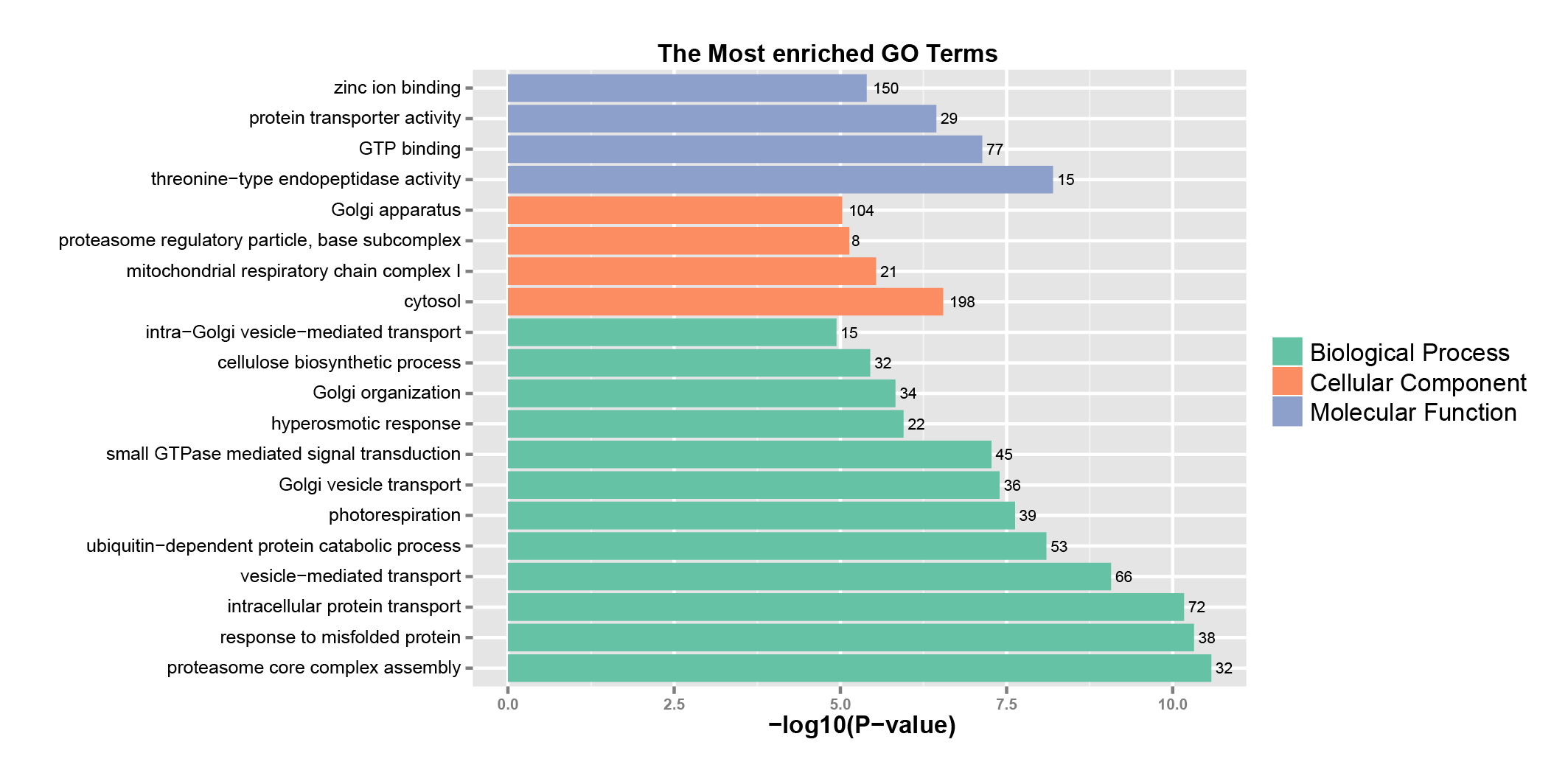
